# The evolution of weaponry and aggressive behaviour in field crickets

**DOI:** 10.1101/2024.12.11.627984

**Authors:** Kevin A. Judge, Briana E. Smith, Shawna L. Ohlmann, Deanna K. Steckler, Alexandria T. Kellington, William H. Cade

## Abstract

Weapons are among the most extravagant sexually selected traits known, yet the evolution of weapon diversity remains understudied. We used field crickets (Orthoptera, Gryllinae) to test two hypotheses explaining interspecific diversity in weaponry. We raised eight species of *Gryllus* field crickets under common garden conditions and staged interactions between conspecific males. We measured body size and weapon shape (relative head and mouthpart size) to determine weapon allometry in both males and females, quantified the intensity of male-male aggression for each species, analyzed the effects of both body size and weapon shape on contest outcome, and tested comparative relationships between morphology and behaviour using phylogenetic least squares regression. We found that larger males won more contests than smaller males in seven of eight species, and weapon shape predicted contest success in only one species. Contrary to the fighting advantage hypothesis, body size was not related to aggressiveness across species, but weaponry was. Additionally, the most aggressive species had the most elaborate weaponry, contrary to the weapon-signal continuum hypothesis. Our results highlight the complexity of weaponry evolution in a group of organisms that has been a model system for the observation and study of aggression for approximately 1000 years.

## Introduction

Sexual selection is a powerful evolutionary force leading to the elaboration of traits that are advantageous in competition for mating opportunities (Darwin, 1871; Andersson, 1994). Most of both theoretical and empirical work on sexual selection has focused on mate choice, both because it was initially more controversial (Cronin, 1991; Shuker & Kvarnemo, 2021), and because the process is more complex than intrasexual contests (McCullough et al., 2016). Understanding of intrasexual selection has also been considered by some to be “essentially complete” since Darwin (Jones & Ratterman, 2009). This bias towards mate choice has left the study of intrasexual selection with major questions that have received scant theoretical or empirical attention (McCullough et al., 2016).

One of the major areas of research facing students of intrasexual selection concerns the evolution of weapon diversity (Emlen, 2008, 2014a; McCullough et al., 2016). Weapons are morphological traits that affect an individual’s success in direct physical contests for access to mates and/or the resources critical to acquiring mates (i.e., “armaments” sensu McCullough & O’Brien, 2022; “dominance traits” sensu Rico-Guevera & Hurme, 2019). Weapons are found in a wide variety of animal taxa (reviewed in Emlen, 2008; but see Tuschhoff & Weins, 2023) and display a bewildering diversity of form (reviewed in Emlen, 2008). Several hypotheses have been proposed to explain the diversity of weaponry within lineages (Emlen, 2008, 2014a; McCullough et al., 2016). In some taxa, weapon diversity appears to be governed by the context in which the competitive sex fights for mates and/or resources used by mates: those that are easily monopolized (e.g., fighting within a burrow) have evolved extreme weapons whereas weapons are reduced or absent when a monopoly access is less likely (e.g., fighting in the open; Emlen & Philips, 2006, reviewed in Emlen, 2014a). In other taxa, performance or fighting advantage during contests, measured as the extent of damage, appears to select for diversity in weapon form (Emberts et al., 2021).

The fighting advantage hypothesis (McCullough et al., 2016), was originally framed as a mechanism by which novel weapon forms (i.e., weapon shape) could be selected because bearers of novel weapons would deliver costs to opponents essentially unopposed (e.g., as the English longbow did against French mounted armour at the Battle of Crécy; Emlen, 2014a). However, fighting advantage can be gained not just through the evolution of new weapon shape, but also increasing weapon size (Darwin, 1871; Andersson, 1994) or both. Separating the effects of weapon shape and weapon size on performance during contests can be challenging, not least because of weapon allometry (McCullough & O’Brien, 2022). However, to the extent that weapon shape and size can be measured, the fighting advantage hypothesis predicts that, if individual variation in weapon size and/or shape predict the outcomes of intraspecific contests, then interspecific variation in the intensity of weapon use should be positively related to interspecific variation in either weapon shape, size or both. We know of no study that has related the performance of weaponry within species to across species variation in weapon size or shape.

Recently, McCullough and O’Brien (2022) proposed that in many species, weapons lie along a continuum of use ranging from “pure weapons” to signals. Pure weapons are those that are used solely in the context of combat and have no utility in signalling to opponents (e.g., stabbing legs in bulb mites) while signals are those whose only function is to signal fighting ability and are never physically engaged in combat (e.g., eye stalks of stalk-eyed flies; reviewed in McCullough & O’Brien, 2022). This weapon-signal continuum makes several predictions. Moving along the continuum from pure weapons to signals should be related to: 1) a decrease in the frequency of the most intense fighting, including fatal fighting, where the weapons are actively engaged in combat, and 2) an increase in the allometric slope of weaponry relative to a reference trait (McCullough & O’Brien, 2022). Thus, the weapon-signal continuum predicts that the allometric slope of weaponry will vary inversely with the frequency of weaponry use. This hypothesis has not yet been tested within a clade of closely related species.

Field crickets (Orthoptera, Gryllidae, Gryllinae) are a model system for the study of aggressive behaviour because they are diverse and distributed globally (Alexander, 1957; Otte & Alexander, 1983; Otte & Cade, 1983; Otte & Perez-Gelabert, 2009; Weissman & Gray, 2019; Cigliano et al., 2023), are easily raised under laboratory conditions (Rakshpal, 1962; Wineriter & Walker, 1988; Judge et al., 2008), and readily display male-male aggression (Chia, 1260; White, 1789; Darwin, 1871; Alexander, 1961; Dixon & Cade, 1986; Adamo & Hoy, 1995). Furthermore, staging and gambling on field cricket fights has been a feature of Chinese culture ever since the Sung Dynasty (AD 960-1278) (Chia, 1260; Laufer, 1927; Jin & Yen, 1998; Suga, 2006). Male field crickets fight aggressively with one another for access to territory and females (Darwin, 1871; Loher & Dambach, 1989) and the largest male usually establishes dominance over smaller rivals (e.g., Alexander, 1961; Dixon & Cade, 1986; Hack, 1997; Shackleton et al., 2005; Kim et al., 2011). Aggressive contests between male field crickets start with low energy and low risk behaviours (e.g., mutual antennal fencing) and, if neither male retreats, can escalate to the highest energy and most risky behaviours where the pair grapple with their mandibles and maxillae (Alexander, 1961; Hack, 1997; Hofmann & Schildberger, 2001; Judge & Bonanno, 2008; Fig. 1). Males have larger heads and mouthparts, including both mandibles and maxillae, than females (Walker et al., 2008; Judge & Bonanno, 2008; Madera & Judge, 2023). Males with proportionately larger heads and mouthparts dominate males with smaller heads and mouthparts – a conclusion known from Chinese cricket-fighting culture (Chia, 1260; Laufer, 1927) – but only in contests that escalate to grappling (Judge & Bonanno, 2008; Judge et al., 2011; but see Briffa, 2008). This is because larger heads and mouthparts increase bite force (Field & Deans, 2001), which is also a predictor of fight success (Hall et al., 2010). There is evidence that intrasexual selection through male-male contest competition has shaped morphological evolution in field crickets. Species of field crickets with relatively narrow heads (Alexander, 1957) never escalate to grappling with their mouthparts in pairwise contests and vice versa (Jang et al., 2008). In other words, species with the smallest weapons are the least aggressive, and species with the largest weapons are most aggressive – a prediction of the fighting advantage hypothesis. Thus, field crickets represent an ideal model system for studying the influence of intrasexual selection on the evolution of interspecific diversity in weaponry.

**Figure 1:**
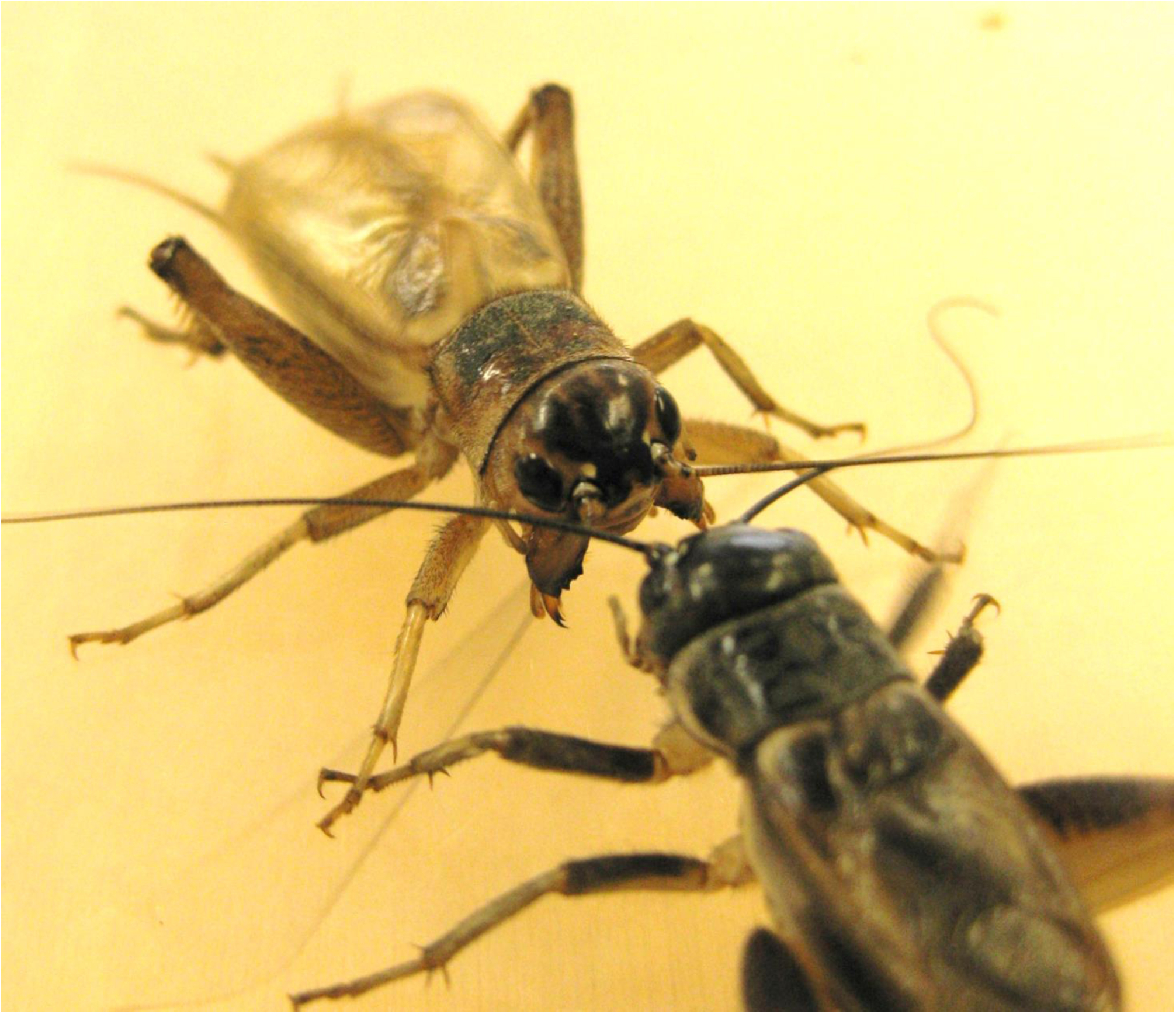
Two adult male variable field crickets (*Gryllus lineaticeps*) engaged in an aggressive interaction. Both males are lashing the opponent with their antennae (note the motion blur), and the facing male is both stridulating (note the motion blur of the raised forewings) and spreading his mandibles and maxillae. The maxillae project below the more robust maxillae and are composed of two distal elements, the pointed lacinia and rounded galea (see also Fig. 1 of Judge & Bonanno, 2008). If neither retreats, then male *Gryllus* field crickets invariably escalate to grappling with their mouthparts.

In the present study, we use a common garden approach with eight species in the North American field cricket genus *Gryllus* to quantify: 1) sexual dimorphism, allometry and interspecific variation in both body size and weapon shape, 2) the effect of variation in both body size and weapon shape on the outcome of aggressive contests between males of the same species, and 3) species-level indices of aggressiveness (i.e., the tendency to escalate to grappling) and morphology. We then use a recently published phylogeny of the genus (Gray et al., 2020) to conduct a comparative analysis of the relationship between aggressiveness and morphology, and test predictions of both the fighting advantage hypothesis (McCullough et al., 2016) as well as the weapon-signal continuum hypothesis (McCullough & O’Brien, 2022).

## Methods

### Study Animals

We established colonies of eight species of North American *Gryllus* field crickets (*G. firmus*, *G. integer*, *G. lineaticeps*, *G. ovisopis*, *G. pennsylvanicus*, *G. rubens*, *G. texensis*, and *G. veletis*). Species were obtained through a combination of field-collecting and soliciting colleagues with existing laboratory colonies (Supplementary Table 1). All experimental animals were raised under the same conditions (Judge et al., 2010), except for where the life history of the species required unique conditions (e.g., Rakshpal, 1962). In brief, adults were housed in large plastic bins (50cm L X 35cm H X 40cm W) containing layers of cardboard egg carton for shelter, glass shell vials filled with water and stoppered with cotton for moisture, and ad libitum pelleted cat chow (Iams Original with Chicken) and rabbit chow (Martin Little Friends) for food. We changed water vials and added food weekly.

When crickets began to enter their last juvenile instar (identified unambiguously by relatively large wing pads) we began to house males individually in plastic deli containers (9 cm diameter, 8 cm high) with a piece of cardboard egg carton for shelter, a microfuge tube filled with water and stoppered with cotton for moisture, and ad libitum cat and rabbit chow. When isolating juvenile males, every effort was made to collect all late instars; this ensured that the available range of male sizes was used (i.e., that we were not biased against including small males). We maintained a pool of between 36 and 72 last instar juvenile males to ensure a steady flow of adult males for aggression trials. Once we had isolated male nymphs, we checked them once a day, except on weekends, to identify newly moulted adults.

We gave newly moulted adult male field crickets an individual identification number and weighed them to the nearest 0.01 mg on an electronic balance (Denver Instruments, model SI-215D). We then randomly paired males that were no more than 5 days different in age. Pairs of males were then scheduled to fight on, or shortly after, the day that the younger male was five days post adult moult because field crickets are mature at this age (Murray & Cade, 1995; KAJ pers. obs.).

### Agonistic Trials

Twenty-four hours prior to a pair’s scheduled trial day, each male was marked with a dot of one of two randomly assigned colours of nail polish. On the day of the trial, we randomly assigned an order to the trials scheduled for the day. Pairs of males were placed into opposite sides of the arena for a minimum of two minutes to acclimate. The arena was a 12.6 cm diameter, 5.5 cm high Plexiglas tube resting on a paper towel. The arena had an opaque plastic divider splitting it into halves with two clear plastic covers (14 cm by 8 cm) over each half of the arena to prevent crickets from jumping out. During acclimation, a dim red lamp provided illumination to the experimenter. We positioned a video camera (SONY Handycam HDR-CX560V) mounted on a tripod directly over the arena. This cameral had infrared vision. The trial was initiated by starting recording and then gently removing the divider, pushing the covers together and immediately leaving the room. Interactions were recorded for 10 minutes and then males were gently separated into individual plastic cups. Each male was then weighed to the nearest 0.01 mg on an electronic balance (Denver Instruments, model SI-215D) and then returned to his individual container. Between trials, all surfaces of the arena that were potentially in contact with crickets were wiped down with 70% ethanol, and the paper towel substrate was replaced.

Environmental conditions in the room used for the trials were the same as conditions for general husbandry given above. We performed all trials during the dark phase of the light cycle, starting approximately 30 minutes after the onset of nighttime. All individuals who took part in trials were held individually for a minimum of two days following their trial and checked to make sure they were still alive and active. If a cricket was found dead (a rare occurrence) it was assumed he had been near death on the day of his trial, so that interaction was not used for analysis. All males were then euthanized by freezing and preserved in 70% ethanol for subsequent measurement of morphology.

### Video Analysis

To minimize any bias during scoring of videos, all videos were assigned a viewing order randomly determined within each species. We then viewed videos in order using the software provided with the video camera (Sony PMB) in blocks of 10 videos alternating among species in a haphazard order. Because our main objective in the present study was to examine the relationship between contest intensity, outcome and morphology, we restricted the behavioural data recorded from each video to a few simple variables. For each video, we recorded if males made physical contact during the 10-minute video recording, and if not, then the trial was excluded from all further analyses. If contact was made, we recorded the identity of the winner, where the winner was defined as the individual from whom the loser retreated in three consecutive physical encounters. Winners also frequently chased the loser, tremulated, and stridulated in a victory display (Hofmann & Schildberger, 2001), but the extent and combination of these behaviours were both variable within and among species whereas retreat by the loser was universal. If no winner was identified by these criteria, the contest was scored as a draw. We also recorded the maximum intensity reached in each contest using a modified version of the intensity scale of Hofmann and Schildberger (2001). Contest intensity values were as follows: 0 = no aggressive behaviours observed, 1 = immediate dominance (i.e., loser and winner apparent upon first physical contact), 2 = antennation by one or both males, 3 = mouthpart gaping, tremulation, and/or kicking by one or both males, and 4 = grappling. Because previous research has shown that weaponry is only important in contests that escalate to grappling (Judge & Bonanno, 2008), we converted contest intensity into a dichotomous variable based on whether the contest escalated to grappling or not. Thus, for each pairwise interaction in which the males contacted one another we have four variables: 1) whether there was a winner or the contest was a draw (Outcome: clear or draw), 2) the identity of the winner in the subset of contests that had a clear outcome (Winner ID), 3) contest intensity (range: 0 to 4), and 4) whether the contest escalated to grappling or not (Grapple: yes or no).

### Measurement of Morphology

Morphological measurement proceeded as in Judge and Bonanno (2008). In brief, all individuals were dissected and positioned in standard orientations and photographed by a digital camera (Infinity Lumenera) attached to a stereomicroscope (Wild M5) using the software Infinity Capture. Specimens were submerged in 70% ethanol during photography to reduce both optical glare and artefacts due to drying out. Measurements were carried out as in Dupuis et al. (2020). Briefly, all individuals had three standardized photographs taken: 1) posterior view of the decapitated head capsule, perpendicular to the transverse axis, 2) dorsal view of the pronotum perpendicular to the transverse axis, and 3) lateral view of both femora placed side by side medial side down. From these three photographs, five morphological dimensions were measured by placing landmarks using the software tpsDig2 (after first creating tps files using tpsUtil; Rohlf, 2016) and then importing the resulting marked tps files into Microsoft Excel and calculating linear dimensions using Pythagoras’ Theorem. The five morphological dimensions were: maxillae span (MS), head width (HW), pronotum length (PL), left femur length (LFL) and right femur length (RFL) (see Judge & Bonanno, 2008, particularly Fig. 1, for a detailed description of each dimension). We calculated mean femur length (MFL) for each individual as the average of LFL and RFL because we detected no evidence of absolute or directional asymmetry (analyses not shown). For each species, we measured both adult males from 90 pairwise contests and a haphazard sample of 90 adult females collected from the breeding bin. Measurements of both males were used in within-species analyses of contest outcomes, however only a subset of 90 males (one chosen randomly from each of 90 pairs) was used for calculation of species-level indices of morphology. This sample size was chosen based on previous studies of sexual dimorphism in morphology (Judge & Bonanno, 2008; Madera & Judge, 2023).

### Calculation of Indices of Body Size and Weapon Shape

We calculated body size as the geometric mean of HW, PL and MFL following the recommendation of Madera and Judge (2023). Geometric mean size (GMS) as an index of body size has an advantage over principal component measures of size in that it is sample independent, thus allowing for comparisons among species, which is a goal of the current study (see Madera & Judge, 2023 for further discussion). We calculated weapon size as the geometric mean of MS and HW and converted this to a shape ratio by dividing by GMS (Mosimann, 1970). Shape ratios have the advantage of being: a) easily understood and directly related to what was physically measured, b) sample independent, and c) measurable for individuals and therefore useful for identification. Furthermore, including shape ratios in statistical analyses utilizes all measured variation, unlike statistical metrics where some fraction of measured variation is discarded (e.g., typically principal components with eigenvalues less than 1). And when both shape ratios and statistical measures of shape give the same results, the use of shape ratios is preferable due to their comparability across studies.

### Statistical Analysis

For within species analyses of the determinants of contest outcome, we treated each contest as the independent experimental unit as in Judge and Bonanno (2008). For each contest, one male was randomly selected to be the focal individual (F) and the other the opponent (O), and then we calculated differences (F-O) in both body size (GMSD = GMSF - GMSO) and weaponry (WSSRD = WSSRF - WSSRO). These differences then became the independent variables in a logistic regression with contest outcome as the binary dependent variable, (0 = focal lost, 1 = focal won). In this way, positive differences in our independent variables mean that the focal male had larger values and was therefore predicted to win, and negative differences mean that the focal male had smaller values and was therefore predicted to lose. To avoid issues with multicollinearity, we conducted stepwise logistic regression. However, because we have clear a priori predictions about the relative importance of our independent variables, we specified their entry into the model in separate blocks in the following order: GMSD and then WSSRD. These analyses were performed on all contests where a clear winner could be identified, as well as only the subset of contests that escalated to grappling because weaponry has previously been shown to be only important during the most intense contests where males grappled with their mouthparts (Judge & Bonanno, 2008).

To assess the allometry of weaponry, we used ordinary least squares regression to estimate the allometric relationship between log weapon size (LGWS) and log geometric mean size (LGGMS) for males and females separately. To test for sexual dimorphism in allometry we conducted a univariate GLM with LGWS as the dependent variable, and LGGMS, Sex and their interaction as independent variables for each species separately. To test for interspecific variation in allometry we conducted a univariate GLM with LGWS as the dependent variable, and LGGMS, Species and their interaction as independent variables. Because of sexual dimorphism in weapon size allometry (see below), separate analyses were conducted for males and females.

If selection due to male-male aggressive contests is important in driving the evolution of male body size and shape in *Gryllus* field crickets, then we predict that factors that consistently affect contest outcome within species will be related to aggressiveness across species, whereas factors that do not affect contest outcome or do so in different directions among species will not be related to aggressiveness across species. Furthermore, to the extent that fecundity selection on females is random with respect to male aggressiveness, we predict that strong selection due to male-male aggression will result in male aggressiveness being related to sexual dimorphism in factors that consistently predict contest outcome within species, but not to sexual dimorphism in factors unrelated or inconsistently related to contest outcome. To examine these predictions, we calculated mean values for both male body size and weaponry, as well as sexual dimorphism (SD: male average/female average) of body size and weaponry. We then conducted phylogenetic generalized least squares (PGLS) regression using the phylogeny of North American Gryllus from Gray et al. (2020; nexus file available in Dryad). PGLS is a variant of generalized least squares regression that accounts for relatedness among taxa by weighting the covariance structure of cross-species data using the evolutionary relationships represented in a phylogeny (Symonds & Blomberg, 2014). The predictor was species’ aggressiveness, measured as the percentage of trials that escalated to grappling (percent grappling), for each of four regression analyses with: mean male body size, mean male weaponry, SD in body size, and SD in weaponry as the response variables.

To test predictions of the weapon-signal continuum, we used PGLS to regress the: a) allometric slope of male weaponry on percent grappling, b) sexual dimorphism in allometry (male slope/female slope) on percent grappling, and c) coefficient of variation in male weaponry on percent grappling. In all cases, the weapon-signal continuum predicts a negative relationship between weapon usage (percent grappling) and each of the dependent variables.

Within species analyses were performed in IBM SPSS Version 28 (IBM Corporation, Armonk, NY). PGLS was performed in R using the packages phytools, nlme, and ape (R Core Team, 2023). All analyses were two-tailed with a Type I error rate of 5%.

## Results

### Summary of Agonistic Contests

We staged and video-recorded 973 unique 10-minute male-male interactions distributed unevenly across eight species of North American *Gryllus* field crickets (Table 1). Of these 973 trials, 13 were discarded because of experimenter error (e.g., video file lost, specimen or id label lost, etc.), 25 were discarded because the males failed to make physical contact during the 10-minute recording, 9 were discarded because one or both males died during the 48 hour period following the trial, and 8 were discarded because one or both males was found to have a physical deformity that was not detected in advance of pairing (e.g., missing tarsus, etc.). This left 918 trials available for analysis, distributed unevenly among the eight species (range = 95 to 129, Table 1). To create a balanced dataset for final analysis we decided to analyze only a random subset of 90 trials per species. Two previous studies detected an effect of weapon size on contest outcome with smaller sample sizes (Judge & Bonanno, 2008, N = 86; Judge et al., 2010, Experiment 1 N = 73, Experiment 2 N = 43) so a sample of 90 trials per species should be adequate to detect biologically meaningful effect sizes.

**Table 1:** A full accounting of the numbers of 10-minute trials staged and recorded on video, along with the numbers of trials excluded from use because of either experimenter error, a lack of contact between males during the 10-minute trial period, one or both males dying during the 48 hours immediately following the trial, or one or both males having a deformity or physical damage that was not noticed prior to pairing. Useable trials represent the difference between recorded trials and the excluded trials. To achieve a balanced design for statistical analysis, we randomly selected 90 trials from the useable for analysis.

| Species | Recorded | Excluded |  |  |  | Useable | Analyzed |
| --- | --- | --- | --- | --- | --- | --- | --- |
|  |  | Error | No Contact | Death | Deformity |  |  |
| *G. firmus* | 132 | 1 | 2 | 0 | 0 | 129 | 90 |
| *G. integer* | 120 | 3 | 3 | 0 | 1 | 113 | 90 |
| *G. lineaticeps* | 100 | 1 | 1 | 0 | 3 | 95 | 90 |
| *G. ovisopis* | 143 | 0 | 13 | 0 | 1 | 129 | 90 |
| *G. pennsylvanicus* | 122 | 1 | 0 | 1 | 1 | 119 | 90 |
| *G. rubens* | 118 | 2 | 1 | 3 | 1 | 111 | 90 |
| *G. texensis* | 129 | 5 | 3 | 5 | 1 | 115 | 90 |
| *G. veletis* | 109 | 0 | 2 | 0 | 0 | 107 | 90 |
| Totals | 973 | 13 | 25 | 9 | 8 | 918 | 720 |

### Contest Intensity

The most common contest intensity level for the experiment was antennation (intensity 2: 255/720, 35.4%), followed by escalation to grappling (intensity 4: 234/720, 32.5%), but this result varied widely among species (Table 2). *G. integer* displayed the highest level of aggression, escalating to grappling in 63.3% of trials, whereas *G. ovisopis* escalated to grappling least frequently at 4.4% (Fig. 2, Table 2). Average contest intensity was highly positively correlated with percentage grappling (r = 0.985, p < 0.001) and so hereafter we only present percent grappling as a measure of species’ aggressiveness because of its direct mechanistic link to our hypotheses concerning the importance of relative weaponry size. Analyses using average contest intensity instead of percent grappling were virtually identical (KAJ et al. unpubl. data).

**Figure 2:**
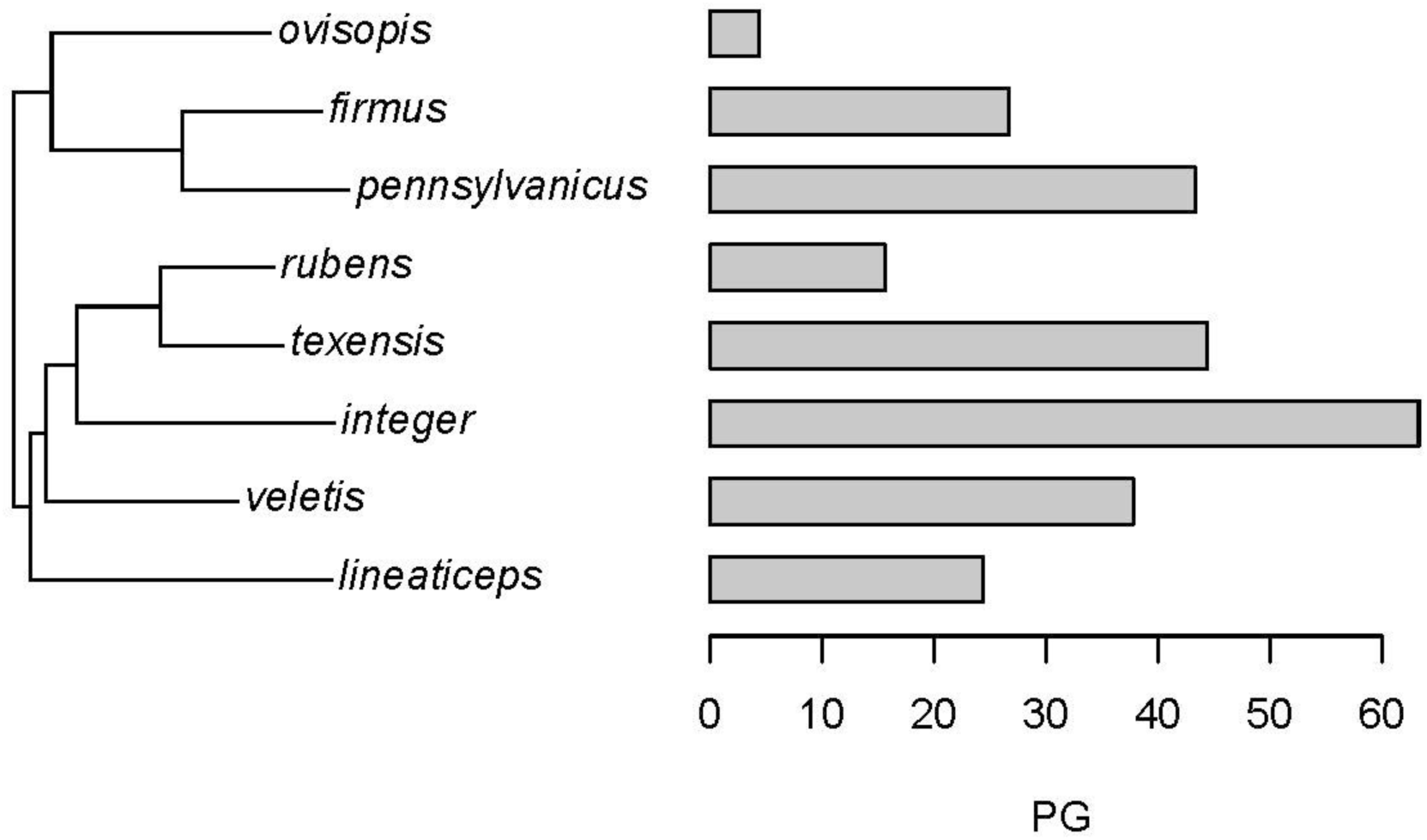
Phylogenetic tree of eight species of North American *Gryllus* field crickets used in this study pruned from the larger tree in Gray et al., 2020. Horizontal bars indicate each species’ aggressiveness as measured by the percentage of trials that escalated to grappling with mouthparts (PG).

**Table 2:** Numbers of trials that escalated to each level of contest intensity, the average contest intensity (ACI) and the percentage of trials that escalated to grappling (PG) for eight species of *Gryllus* field crickets. Contest intensity levels are as follows: 0 = no aggression, 1 = immediate dominance, 2 = antennation, 3 = gaping, tremulation, kicking, and 4 = grappling (see Methods: Video Analysis for details); numbers in parentheses represent the subset of trials where a clear winner could not be determined. N = 90 trials for each species.

| Species | Contest Intensity |  |  |  |  | ACI | PG |
| --- | --- | --- | --- | --- | --- | --- | --- |
|  | 0 | 1 | 2 | 3 | 4 |  |  |
| *G. firmus* | 0 | 1 | 42 | 23(1) | 24 | 2.8 | 26.7 |
| *G. integer* | 2 | 3 | 9 | 19 | 57 | 3.4 | 63.3 |
| *G. lineaticeps* | 1 | 4 | 34(2) | 29 | 22 | 2.7 | 24.4 |
| *G. ovisopis* | 11 | 0 | 29(5) | 46(17) | 4 | 2.4 | 4.4 |
| *G. pennsylvanicus* | 0 | 0 | 27 | 24 | 39 | 3.1 | 43.3 |
| *G. rubens* | 1 | 1 | 59(1) | 15(1) | 14 | 2.4 | 15.6 |
| *G. texensis* | 0 | 3 | 34 | 13 | 40 | 3.0 | 44.4 |
| *G. veletis* | 2 | 7 | 21(1) | 26(1) | 34 | 2.9 | 37.8 |
| Totals* | 17 | 19 | 255(9) | 195(20) | 234 | 2.8 | 32.5 |

\* Values are averages for ACI and PG

### Sexual Dimorphism in Body Size and Weapon Shape

To test for sexual size dimorphism (SSD), we conducted univariate GLMs separately for each species of *Gryllus*, with geometric mean size (GMS) as the dependent variable and SEX as the independent variable. We detected SSD in seven of the eight species, but not in a consistent direction (Fig. 3). In *G. integer, G. lineaticeps, G. texensis* and *G. veletis* males were larger than females (all p < 0.001), whereas in *G. ovisopis, G. pennsylvanicus* and *G. rubens* females were larger than males (all p < 0.030); we failed to detect SSD in *G. firmus* (p = 0.608, Fig. 3). Analysis of sexual dimorphism in weaponry yielded a consistent pattern across species. Univariate GLMs with weapon size shape ratio (WSSR) as the dependent variable and SEX as the independent variable detected male-biased sexual dimorphism in all eight species (all p < 0.001, Fig. 4).

**Figure 3:**
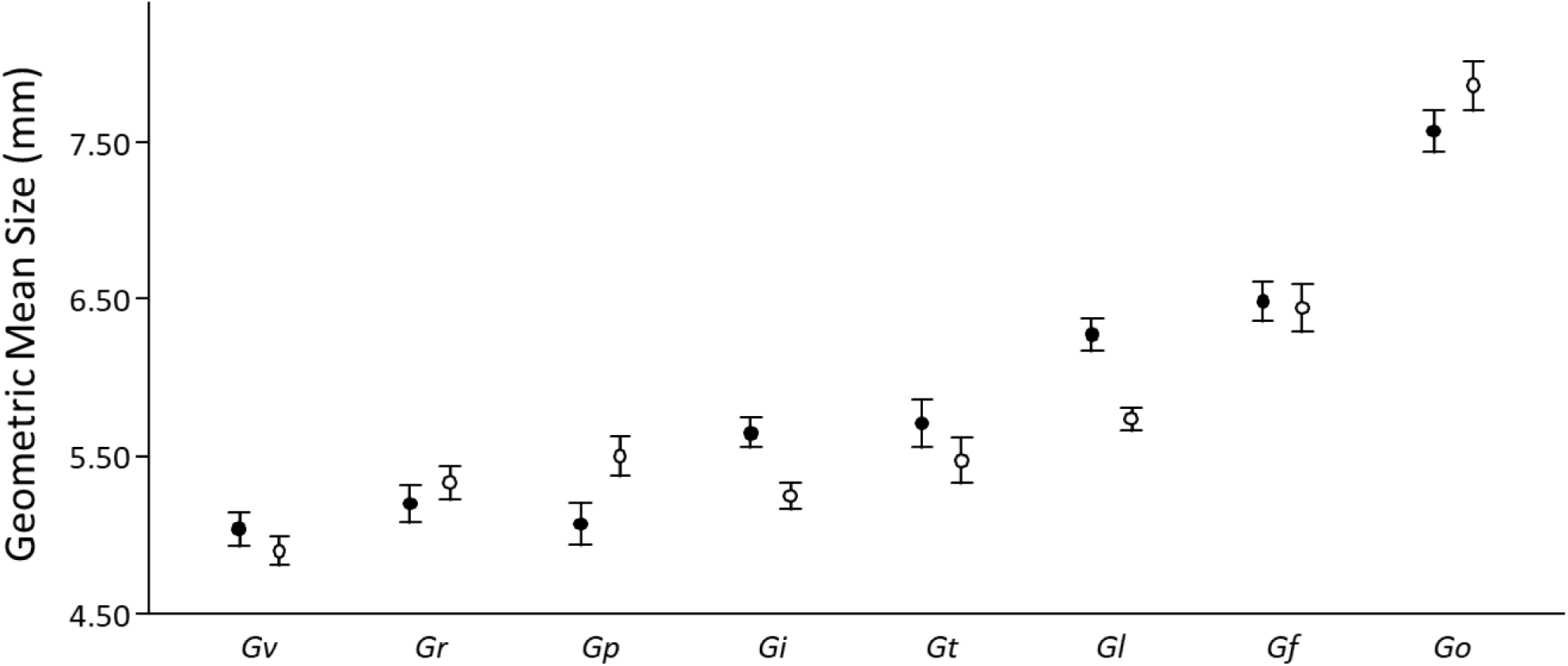
Average geometric mean size (GMS) for adult males (filled circles) and adult females (open circles) of nine species of North American field crickets (*Gf*: *Gryllus firmus*; *Gi*: *G. integer*, *Gl*: *G. lineaticeps*; *Go*: *G. ovisopis*; *Gp*: *G. pennsylvanicus*; *Gr*: *G. rubens*; *Gt*: *G. texensis*; *Gv*: *G. veletis*) arranged in order of increasing average GMS. Error bars are 99% confidence limits.

**Figure 4:**
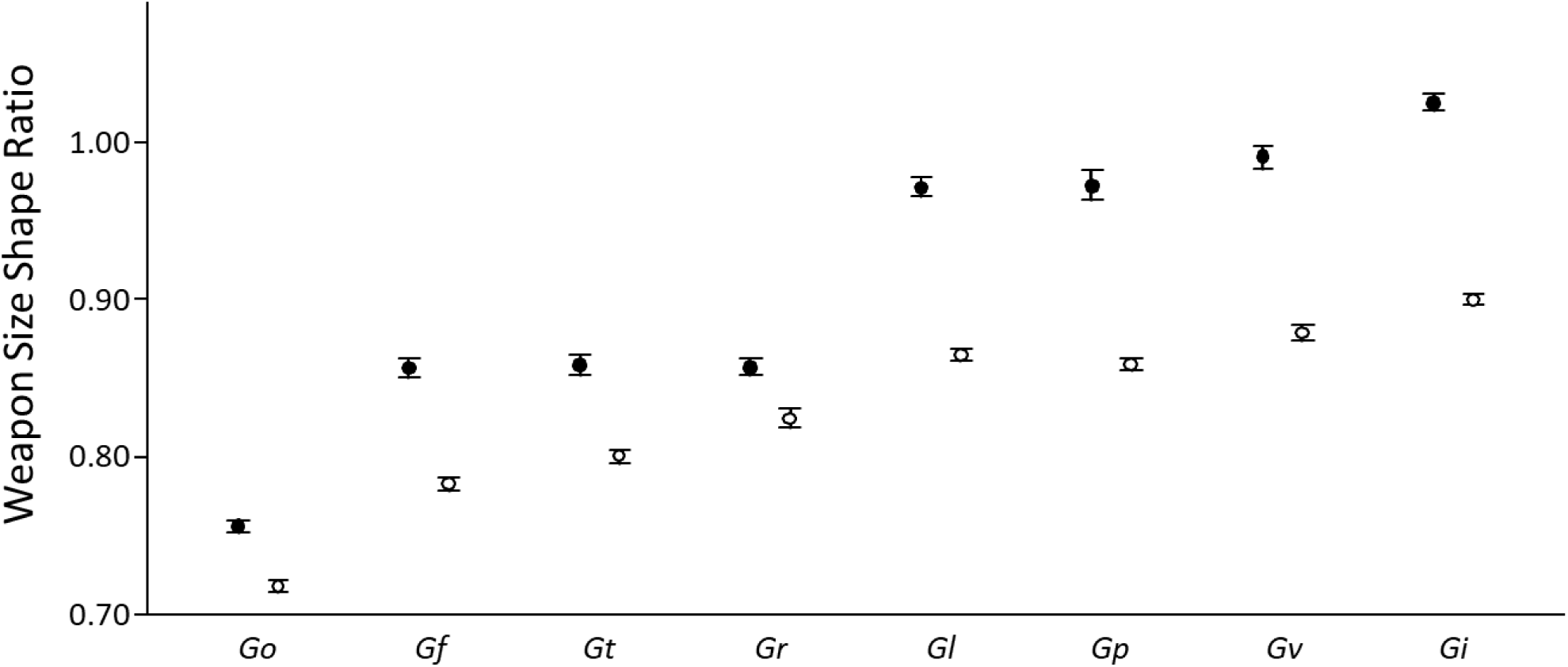
Average weapon size shape ratio (WSSR) for adult males (filled circles) and adult females (open circles) of nine species of North American field crickets (*Gf*: *Gryllus firmus*; *Gi*: *G. integer*, *Gl*: *G. lineaticeps*; *Go*: *G. ovisopis*; *Gp*: *G. pennsylvanicus*; *Gr*: *G. rubens*; *Gt*: *G. texensis*; *Gv*: *G. veletis*) arranged in order of increasing average WSSR. Error bars are 99% confidence limits.

### Within Species Effects of Body Size and Weapon Shape

When considering all contests where there was a clear winner, regardless of contest intensity, body size difference was the only consistent predictor of contest outcome in field crickets. As predicted, in all species the focal male won contests more often when he was larger than his opponent than when he was smaller. This effect was more pronounced the greater the difference in body size between competitors, and statistically significant in all species except *G. ovisopis* (p = 0.353, all other p < 0.001, Table 3). For the seven species where body size advantage significantly predicted winning, body size difference explained between 16.1 and 38.2% of the variation in contest outcome (Table 3). In only one species, *G. integer*, did difference in weaponry improve prediction of contest outcome enough to be entered as a predictor in the second block of the stepwise logistic regression. When focal male *G. integer* had proportionately larger weaponry than their opponent, they were more likely to win than when they had smaller weaponry, and this effect increased with increasing difference in weaponry size (Table 3).

**Table 3:** Summary of stepwise logistic regression results examining the effect of body size (GMS, entered in Block 1) and differences in weapon size shape ratio (WSSR, entered in Block 2) for all trials with a clear winner. Nagelkerke R^2^ is a measure of strength of association and B is the logistic slope value for either body size or weapon size shape ratio. Bold rows represent steps that are statistically significant at p < 0.05.

| Species | N | Block | $\chi^2$ | df | p | Nagelkerke R<sup>2</sup> | B(GMS) | B(WSSR) |
| --- | --- | --- | --- | --- | --- | --- | --- | --- |
| *G. firmus* | 89 | **1** | **14.277** | **1** | **<0.001** | **0.198** | **1.523** |  |
|  |  | 2 | 1.470 | 1 | 0.225 | 0.216 | 1.324 | 11.365 |
| *G. integer* | 88 | **1** | **21.659** | **1** | **<0.001** | **0.291** | **2.666** |  |
|  |  | **2** | **4.247** | **1** | **0.039** | **0.340** | **2.010** | **21.389** |
| *G. lineaticeps* | 87 | **1** | **12.193** | **1** | **<0.001** | **0.175** | **2.073** |  |
|  |  | 2 | 2.780 | 1 | 0.095 | 0.212 | 2.552 | -16.935 |
| *G. ovisopis* | 57 | 1 | 0.861 | 1 | 0.353 | 0.020 | 0.448 |  |
|  |  | 2 | 0.024 | 1 | 0.878 | 0.021 | 0.456 | 2.410 |
| *G. pennsylvanicus* | 90 | **1** | **11.607** | **1** | **<0.001** | **0.161** | **1.589** |  |
|  |  | 2 | 0.702 | 1 | 0.402 | 0.171 | 1.320 | 6.051 |
| *G. rubens* | 87 | **1** | **29.148** | **1** | **<0.001** | **0.382** | **2.731** |  |
|  |  | 2 | 1.122 | 1 | 0.289 | 0.394 | 2.965 | 10.008 |
| *G. texensis* | 90 | **1** | **12.211** | **1** | **<0.001** | **0.169** | **1.187** |  |
|  |  | 2 | 0.448 | 1 | 0.503 | 0.175 | 1.279 | -5.949 |
| *G. veletis* | 86 | **1** | **15.428** | **1** | **<0.001** | **0.219** | **1.852** |  |
|  |  | 2 | 0.368 | 1 | 0.544 | 0.224 | 1.743 | 5.111 |

Because weaponry has been shown to be important primarily in contests that escalate to grappling with mouthparts (Judge & Bonanno, 2008), we restricted our analyses of contest outcome to only those contests that escalated to grappling (see Table 2). Like the analysis of trials of all intensities, body size difference was a significant predictor of fight outcome in all species except for *G. ovisopis*, and weapon size difference was a significant predictor in only *G. integer* (Table 4).

**Table 4:** Summary of stepwise logistic regression results examining the effect of differences in body size (GMS, entered in Block 1) and differences in weapon size shape ratio (WSSR, entered in Block 2) for all trials with a clear winner that escalated to grappling. Nagelkerke R^2^ is a measure of strength of association and B is the logistic slope value for either body size or weapon size shape ratio. Bold rows represent steps that are statistically significant at p < 0.05.

| Species | N | Block | $\chi^2$ | df | Sig. | Nagelkerke R<sup>2</sup> | B(GMS) | B(WSSR) |
| --- | --- | --- | --- | --- | --- | --- | --- | --- |
| *G. firmus* | 24 | **1** | **8.121** | **1** | **0.004** | **0.386** | **4.012** |  |
|  |  | 2 | 0.687 | 1 | 0.407 | 0.413 | 3.888 | 15.298 |
| *G. integer* | 57 | **1** | **7.442** | **1** | **0.006** | **0.163** | **2.097** |  |
|  |  | **2** | **3.537** | **1** | **0.060** | **0.234** | **1.458** | **21.658** |
| *G. lineaticeps* | 22 | **1** | **5.528** | **1** | **0.019** | **0.296** | **3.976** |  |
|  |  | 2 | 1.571 | 1 | 0.210 | 0.368 | 5.132 | -24.814 |
| *G. ovisopis* | 4 | 1 | 0.235 | 1 | 0.628 | 0.085 | 1.106 |  |
|  |  | 2 | 4.263 | 1 | 0.039 | 1.000 | * |  |
| *G. pennsylvanicus* | 39 | **1** | **10.328** | **1** | **0.001** | **0.312** | **2.416** |  |
|  |  | 2 | 0.461 | 1 | 0.497 | 0.324 | 1.980 | 8.719 |
| *G. rubens* | 14 | **1** | **3.386** | **1** | **0.066** | **0.288** | **2.559** |  |
|  |  | 2 | 3.502 | 1 | 0.061 | 0.522 | 6.087 | 58.174 |
| *G. texensis* | 40 | **1** | **10.445** | **1** | **0.001** | **0.307** | **2.341** |  |
|  |  | 2 | 0.066 | 1 | 0.797 | 0.308 | 2.369 | -3.303 |
| *G. veletis* | 34 | **1** | **10.175** | **1** | **0.001** | **0.352** | **3.598** |  |
|  |  | 2 | 1.747 | 1 | 0.186 | 0.402 | 3.715 | 21.118 |

\* The second block of the logistic model for *G. ovisopis* was over specified due to extremely low sample size of trials that escalated to grappling.

### Allometry of Weaponry

In all eight species of *Gryllus* field crickets, male weaponry was positively allometric relative to female weaponry (SEX * LGGMS interaction: all p < 0.028). And within each sex, the allometry of weaponry varied among species (SPECIES * LGGMS interaction: Males: F_8,792_ = 18.753, p < 0.001; Females: F_8,792_ = 6.816, p < 0.001: Table 5).

**Table 5:**
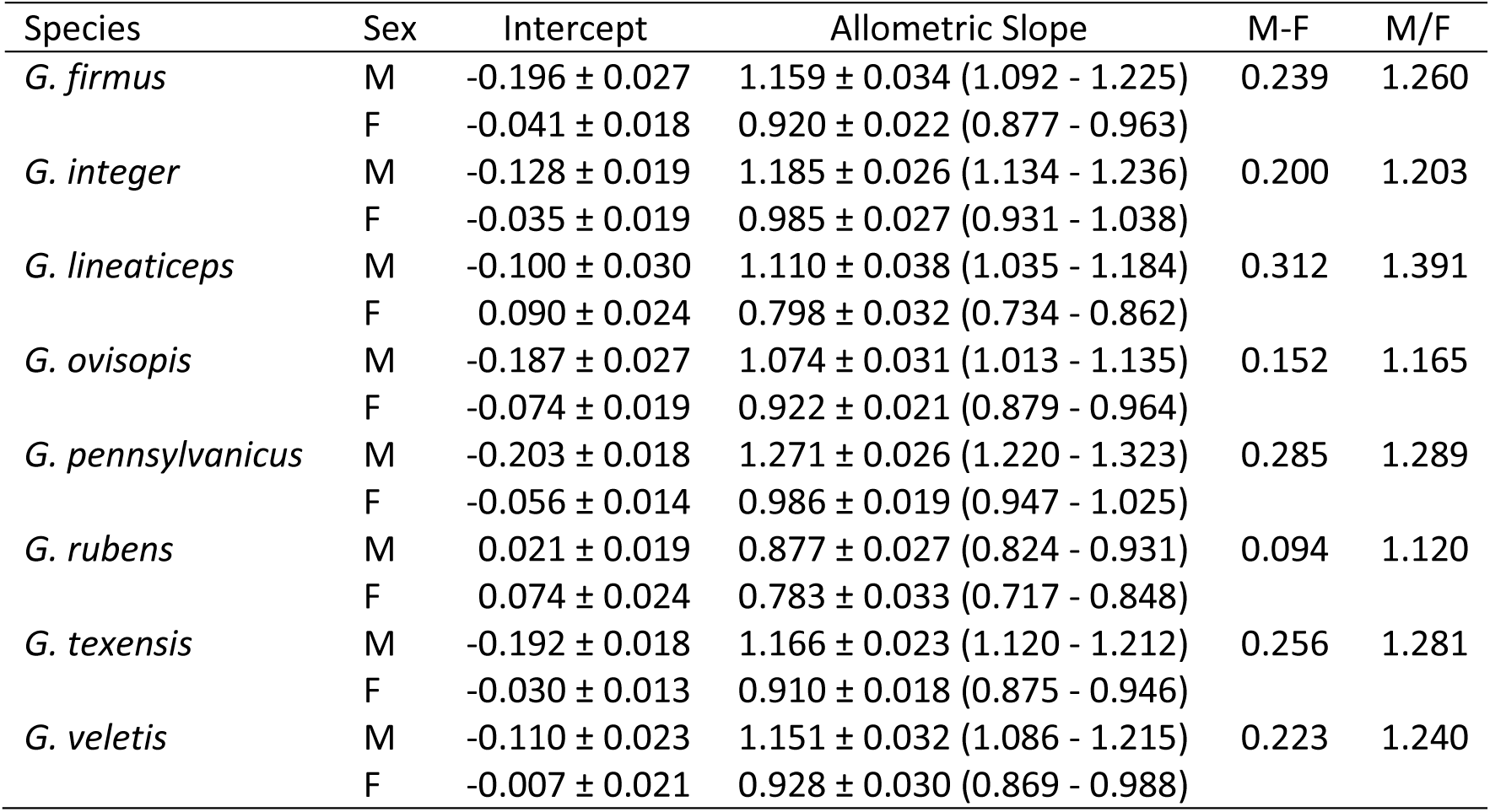
Allometric slopes for ordinary least squares regressions of Log Weaponry Size vs Log GMS for males and females of all eight species of *Gryllus* field crickets. N = 90 for both males and females of every species, ± indicates the standard error of the mean, and values in parentheses are the 95% confidence limits. Sexual dimorphism in allometric slope is calculated as both the difference between male and female slopes (M-F) and their ratio (M/F).

| Species | Sex | Intercept | Allometric Slope | M-F | M/F |
| --- | --- | --- | --- | --- | --- |
| *G. firmus* | M | -0.196 ± 0.027 | 1.159 ± 0.034 (1.092 - 1.225) | 0.239 | 1.260 |
| | F | -0.041 ± 0.018 | 0.920 ± 0.022 (0.877 - 0.963) | | |
| *G. integer* | M | -0.128 ± 0.019 | 1.185 ± 0.026 (1.134 - 1.236) | 0.200 | 1.203 |
| | F | -0.035 ± 0.019 | 0.985 ± 0.027 (0.931 - 1.038) | | |
| *G. lineaticeps* | M | -0.100 ± 0.030 | 1.110 ± 0.038 (1.035 - 1.184) | 0.312 | 1.391 |
| | F | 0.090 ± 0.024 | 0.798 ± 0.032 (0.734 - 0.862) | | |
| *G. ovisopis* | M | -0.187 ± 0.027 | 1.074 ± 0.031 (1.013 - 1.135) | 0.152 | 1.165 |
| | F | -0.074 ± 0.019 | 0.922 ± 0.021 (0.879 - 0.964) | | |
| *G. pennsylvanicus* | M | -0.203 ± 0.018 | 1.271 ± 0.026 (1.220 - 1.323) | 0.285 | 1.289 |
| | F | -0.056 ± 0.014 | 0.986 ± 0.019 (0.947 - 1.025) | | |
| *G. rubens* | M | 0.021 ± 0.019 | 0.877 ± 0.027 (0.824 - 0.931) | 0.094 | 1.120 |
| | F | 0.074 ± 0.024 | 0.783 ± 0.033 (0.717 - 0.848) | | |
| *G. texensis* | M | -0.192 ± 0.018 | 1.166 ± 0.023 (1.120 - 1.212) | 0.256 | 1.281 |
| | F | -0.030 ± 0.013 | 0.910 ± 0.018 (0.875 - 0.946) | | |
| *G. veletis* | M | -0.110 ± 0.023 | 1.151 ± 0.032 (1.086 - 1.215) | 0.223 | 1.240 |
| | F | -0.007 ± 0.021 | 0.928 ± 0.030 (0.869 - 0.988) | | |

### Among Species Relationships

Based on the results of within species analyses, we predicted that, at the species level, aggressiveness would be positively related to male body size and sexual dimorphism (SD) in body size, but not to male weaponry or SD in weaponry. In fact, we found the opposite pattern. Aggressiveness was not related to male body size (t = - 1.288, p = 0.245; Fig. 5a) or SD in body size (t = 0.909, p = 0.399; Fig. 5c), whereas it was positively related to male weaponry (t = 2.582, p = 0.042; Fig. 5b) and SD in weaponry (t = 3.107, p = 0.021; Fig. 5d).

**Figure 5:**
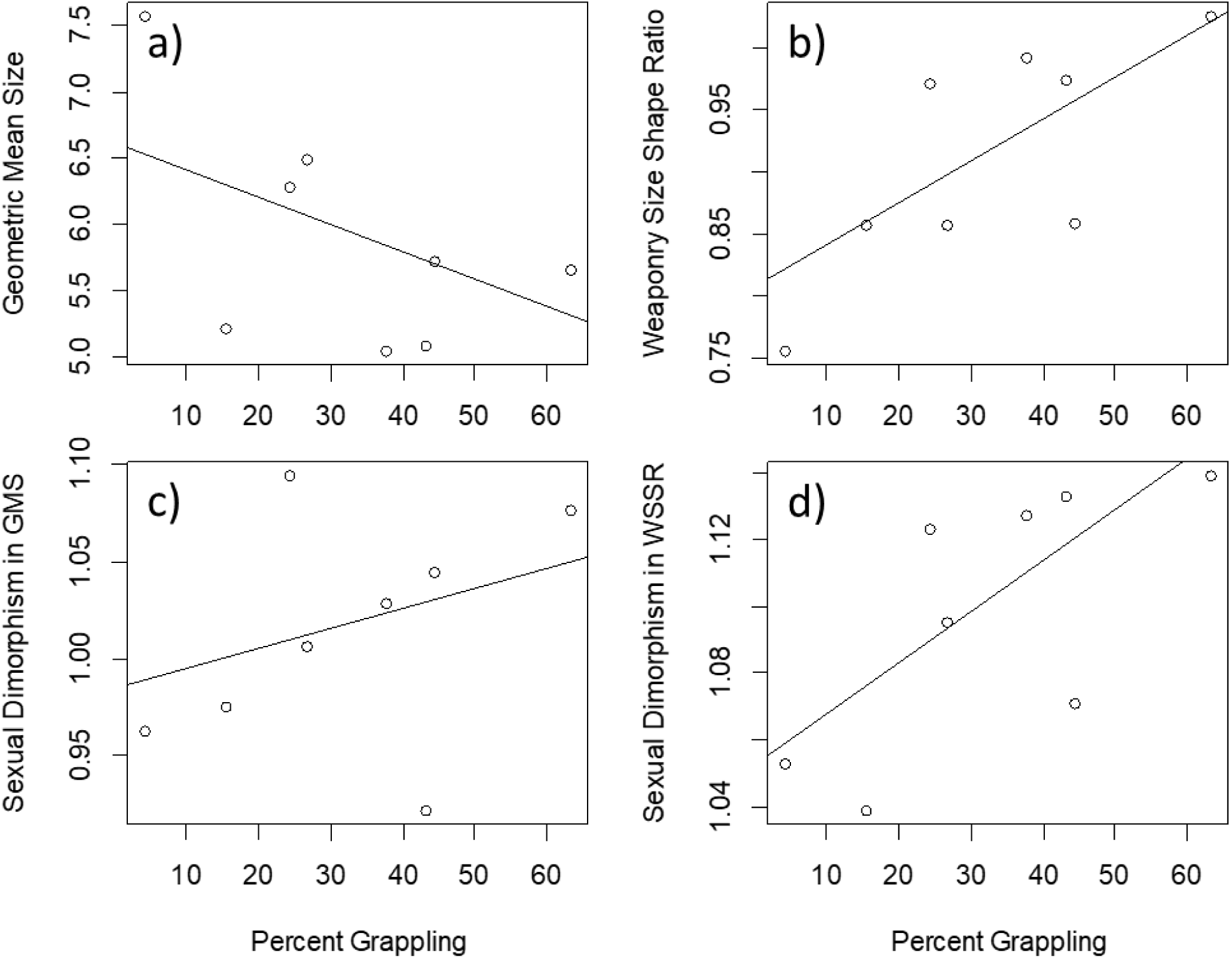
Average: a) male geometric mean size (GMS), b) male weapon size shape ratio (WSSR), c) sexual dimorphism in GMS (GMSSD), and d) sexual dimorphism in WSSR (WSSRSD) versus the percentage of trials that escalated to grappling (PG) for eight species of North American *Gryllus* field crickets. Sexual dimorphism is calculated as average male trait value/average female trait value. Fitted lines are the result of phylogenetic generalized least squares (PGLS) regressions (a: GMS = -0.021PG +6.625, p = 0.245; b: WSSR = 0.003PG + 0.807, p = 0.042; c: GMSSD = 0.001PG + 0.985, p = 0.399; d: WSSRSD = 0.002PG + 1.052, p = 0.021).

Contrary to the predictions of the weapon-signal continuum, allometric slopes, sexual dimorphism in weapon allometry and variation in male weaponry was not negatively related to the use of weaponry across species. The allometric slope of male weaponry was positively related to weapon use (percent grappling) across species (t = 3.156, p = 0.020; Fig. 6a). We were unable to detect any linear relationship between either sexual dimorphism in weapon allometry (t = 1.139, p = 0.298; Fig. 6b) or variation in male weaponry (t = 1.727, p = 0.135; Fig. 6c) and weapon use.

**Figure 6:**
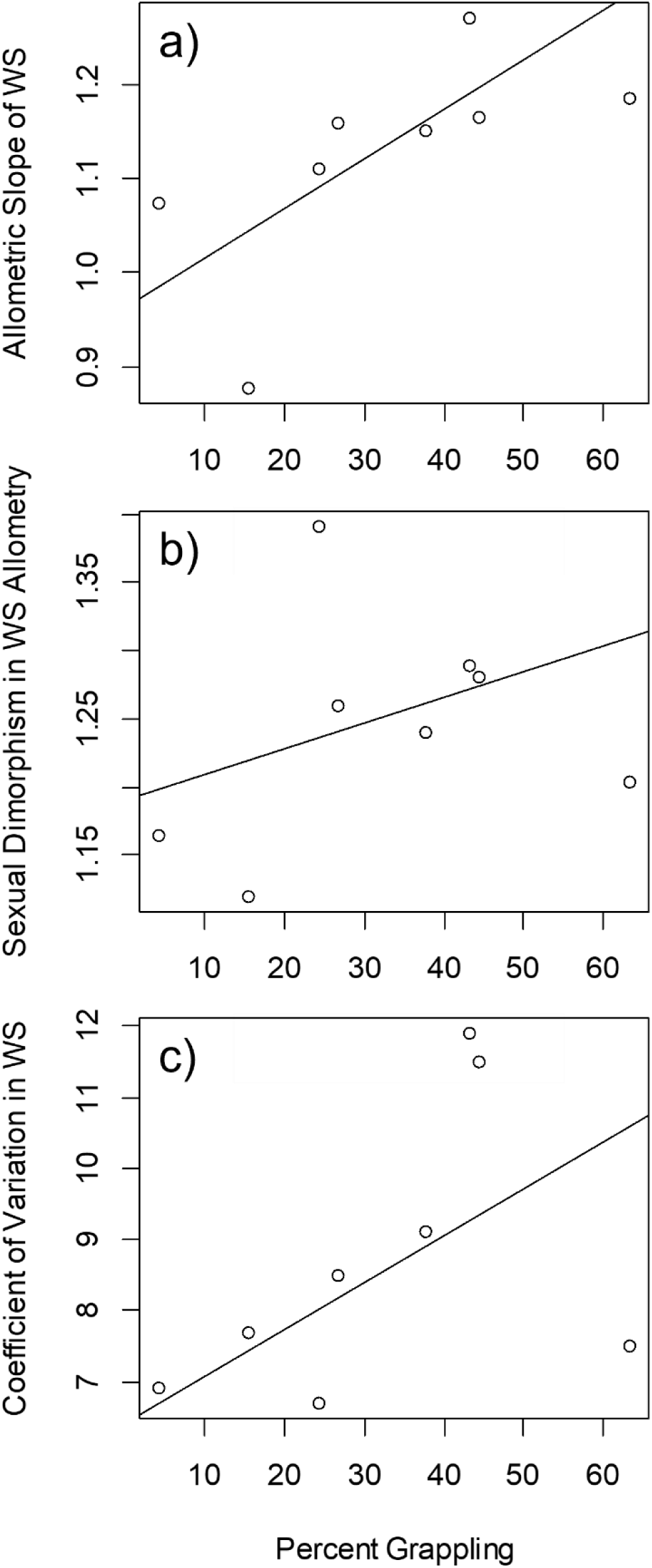
Average: a) allometric slope of male weapon size (WSALLO), b) sexual dimorphism in weapon size allometry (WSALLOSD), and c) coefficient of variation in male weapon size (WSCV) versus the percentage of trials that escalated to grappling (PG) for eight species of North American *Gryllus* field crickets. Sexual dimorphism is calculated as average male trait value/average female trait value. Fitted lines are the result of phylogenetic generalized least squares (PGLS) regressions (a: WSALLO = 0.005PG + 0.961, p = 0.0.020; b: WSALLOSD = 0.002PG + 1.190, p = 0.298; c: WSCV = 0.066PG + 6.387, p = 0.135).

## Discussion

In this study we take a common garden experimental approach to study aggressive behaviour and morphology in eight species of *Gryllus* field crickets. We use the results of within-species analyses of the effects of body size and weaponry to make predictions about what morphological traits should have among-species relationships with aggressive behaviour, and we test predictions of both the fighting advantage and weapon-signal continuum hypotheses. Most *Gryllus* species were sexually size dimorphic, although not in a consistent direction (Fig. 3), whereas all eight species showed male-biased sexual dimorphism in weaponry (Fig. 4). In seven of eight species, larger males won more contests and the magnitude of the advantage increased with greater disparity in size; however, in only one species did weaponry have a significant positive effect on contest outcome (Tables 3 and 4). These results suggest that male-male competition exerts consistent directional selection on male body size but not on male weaponry. Consequently, we predicted that interspecific variation in the level of male aggression would predict interspecific variation in both male body size, and - to the extent that fecundity selection on females is random with respect to male aggression – sexual dimorphism in body size, and not male weaponry and sexual dimorphism in weaponry. However, we detected the opposite pattern. In phylogenetically controlled analyses, interspecific aggression predicted male weaponry and sexual dimorphism in weaponry, but not male body size or sexual dimorphism in body size (Fig. 5).

McCullough and O’Brien (2022) propose that traits that are used during male-male competition (i.e., armaments in their terminology) fall along a continuum from those that are solely used during combat and have little to no function as signals (i.e., pure weapons) to those that function mainly as signals of resource holding potential (RHP; Parker, 1974) to deter rivals as well as indicators of breeding value for potential mates but which may occasionally be used during combat (i.e., pure signals). The weapon-signal continuum predicts that pure weapons will be: negatively allometric, less sexually dimorphic, less variable, and used often in combat that often results in death. Whereas pure signals will be: positively allometric, more sexually dimorphic, more variable, and used for combat rarely and lethal combat more rarely still (McCullough & O’Brien, 2022). Thus, the weapon-signal continuum predicts a negative relationship between weapon use and: a) weapon allometric slope, b) sexual dimorphism in weapon allometric slope, and c) weapon variability. It also predicts a positive relationship between weapon use and lethal fights. The results of our study of *Gryllus* field crickets fail to support any of these predictions as there were positive relationships between weapon use (i.e., percent grappling) and weaponry allometric slope, sexual dimorphism in allometric slope, and weapon variability (Fig. 6) although only the first was statistically significant. The frequency of lethal fighting was also low to the point of undetectability. That interspecific variation in *Gryllus* field cricket weaponry fails to conform to the weapon signal continuum is perhaps unsurprising given that aggressive interactions usually occur at night and/or in burrows where the potential for weapons to play a signalling role is low. It is possible that the weapon signal continuum explains diversity in weapon form at broader evolutionary scales (i.e., among classes, e.g., McCullough & O’Brien, 2022) than among species within a genus. However, there are a several other hypotheses to explain interspecific diversity in weaponry (McCullough et al., 2016) and the lack of conformity of our results with one of those hypotheses simply drives home the point that the evolution of weaponry is deserving of attention from evolutionary biologists (Emlen, 2008, 2014a; McCullough et al., 2016).

Another hypothesis proposed as an explanation of weapon diversity concerns the context in which contests occur, specifically that diversity in the conditions in which fights take place (e.g., within a burrow or out in the open) drives diversity in the form of weaponry (Emlen, 2008, 2014a, b; McCullough et al., 2016). Context has been suggested to play a role in the evolution of aggressiveness in field crickets (Jang et al., 2008; Bertram et al., 2011). Some species of *Gryllus* field crickets primarily inhabit forest and are not known to burrow (Walker, 1974, 2001), and are consequently known by the common name of “wood crickets”, whereas the majority of *Gryllus* species utilize burrows, which males defend aggressively (Loher & Dambach, 1989; Bertram et al., 2011; Rodríguez-Muñoz et al., 2010). The wood crickets, *G. fultoni* and *G. vernalis*, did not escalate to grappling in pairwise contests, and this low level of aggression was attributed to the absence of burrow use (Jang et al., 2008). A more formal phylogenetic analysis linked burrow use with escalation to grappling, but assessed both behaviours as simply present or absent (Bertram et al., 2011). The two wood crickets studied so far are also known to have relatively narrower heads (i.e., smaller weaponry) than species that are known to escalate to grappling (see Fig. 1 in Alexander, 1957) suggesting that fighting in or around the narrow confines of a burrow selects for increased aggressiveness and weaponry, whereas fighting on the open forest floor exerts either weak positive or negative selection for aggressiveness and weaponry (Emlen, 2014a,b). Our results provide partial support for this hypothesis because the least aggressive species in our analysis, *G. ovisopis*, had the smallest and least sexually dimorphic weaponry, and is a wood cricket (Walker, 1974). *G. ovisopis* did still occasionally escalate to grappling, and there was a wide range of variation in both weaponry and aggressiveness amongst the seven other species in our study that are known to use burrows. To explain these results, the context specific hypothesis (Emlen, 2008, 2014a; McCullough et al., 2016) would predict that burrow use is a continuous rather than dichotomous behaviour. We were unsuccessful in developing a burrowing behavioural assay to test this hypothesis, but further efforts may be able to make progress. Alternatively, our results suggest that factors underlying most of the variation in both aggressive behaviour and weaponry are as yet unexplained.

Weapon diversity may be selected for if novel or more elaborate weapons provide an advantage over rivals during aggressive contests (i.e., fighting advantage hypothesis: McCullough et al., 2016). In coreid bugs, males fight over access to females by squeezing rivals with their hind legs which are armed with varying numbers and patterns of sharp spines that deliver damage in the form of puncture wounds (Emberts et al., 2021). Amount of damage depended on hind leg weaponry, and divergent kinds of weaponry were found to deliver similar amounts of damage, suggesting that performance during aggressive contests between male bugs selects for elaboration and diversification of hind leg weaponry (Emberts et al., 2021). Although we failed to detect a weapon size advantage in seven of eight *Gryllus* field crickets, there was an advantage of relatively larger heads and mouthparts in the most aggressive of the eight species, and positive comparative relationships between both weaponry elaboration and sexual dimorphism in weaponry and species’ tendency to use weaponry during escalated contests (Fig. 5). Given that bite force is positively correlated with success in aggressive contests (Hall et al., 2010) and larger weaponry means greater bite forces (Field & Deans, 2001), our results provide some support for the fighting-advantage hypothesis as an explanation of interspecific variation in weaponry in *Gryllus*.

Although interspecific variation in weaponry is related to species’ use of weaponry in aggressive contests, the performance advantage of large weaponry appears to be weak relative to larger body size (this study; see also Briffa 2008) and is more detectable when body size is experimentally controlled (Judge & Bonanno, 2008; Judge et al., 2010; Kelly & L’Heureux, 2021). Despite this large disparity in the effects of body size and weaponry on contest outcome within species, our results suggest that intrasexual selection has had a greater effect on the evolution of weaponry shape than body size in *Gryllus* field crickets. In fact, the pattern of intraspecific variation in body size is the reverse of what we would predict based on sexual selection via male-male contest competition (Fig. 5a). Large body size is favoured by fecundity selection on females in many insects (Honĕk, 1993) and it is possible that the pattern of body size variation in *Gryllus* is the result of strong fecundity selection acting in direct opposition to, and overwhelming, intrasexual selection. We know of no studies that have estimated the relative effects on species’ average body size of both fecundity selection on females and sexual selection on males, however, this hypothesis implies a strong intersexual genetic correlation constraining the ability of the sexes to achieve independent body size optima. Although there is some evidence for such genetic constraint on intersexual variation in optimal body size (Bedhomme & Chippindale, 2007; Poissant et al., 2010), our results suggest that there is some room for sexual selection to shape body size as there was a positive relationship between species’ aggressiveness in male contests and both the degree and direction of sexual dimorphism in body size (Fig. 5c).

Recently, attention has been drawn to the observation that sexually selected traits, and weapons in particular, are apparently absent from most animal phyla, and extremely patchily distributed in the two phyla – Chordata and Arthropoda – that contain the majority of published examples (Tuschhoff & Weins, 2023). Neither asexuality, hermaphroditism, a parasitic lifestyle nor a marine broadcast spawning existence explained the distribution of sexually selected traits across the Animal Kingdom or within Chordata and Arthropoda because, although a great many taxa with those characteristics apparently lack sexually selected traits, so too do a great many species that are sexual, monoecious, free-living and terrestrially-copulating (Tuschhoff & Weins, 2023). Field cricket weaponry is not extreme in comparison to well-known and well-studied taxa that are models for the study of intrasexually selected weaponry (reviewed in Emlen, 2008, 2014a; Rico-Guevara & Hurme, 2019) and has been ignored by western science until recently (Briffa, 2008; Judge & Bonanno, 2008). Practitioners of Chinese cricket fighting have known for over 800 years that male weaponry is an important factor affecting contest competition (Chia, 1260; Laufer, 1927; Suga, 2006). Our results suggest that small variations in body shape can substantially affect macroevolutionary patterns of trait variation within a common and globally distributed group. We ask, how would conclusions regarding the evolution of sexually selected traits change if the vast majority of sexual, monoecious, free-living and terrestrial taxa in fact do possess sexually selected traits? An answer will not come without broader taxonomic sampling and closer examination of apparently well-known organisms.

## Author Contributions

KAJ, BES and WHC conceived of and designed the study; KAJ and BES reared the study animals and conducted the behaviour experiments; KAJ scored the behaviour trials; SO, DS and AK dissected, photographed and measured specimens; KAJ analyzed the data and wrote the initial draft; all authors contributed to editing the manuscript; KAJ and WHC secured funding.

## Acknowledgements

Thanks to William E. Wagner Jr. and Raine Kortet for providing crickets to start lab colonies for two species. Thanks to David A. Gray for help with R coding. Ife Abiola helped with colony maintenance. Thanks to Erin Gaydosh for assistance with cricket measurements. Erin L. McCullough and Darryl T. Gwynne provided helpful comments on an earlier draft of this manuscript. This research was funded by Natural Science and Engineering Research Council Discovery Grants to KAJ (RGPIN-2017-04674) and WHC (RGPIN-2007-06174).

## Data Availability

R code, behavioural and morphological data, and the photographs from which morphological data was measured are available on Dryad. The complete set of contest videos are available on Harvard Dataverse, and all experimental subjects are preserved in 70% ethanol and deposited in the entomology collection at MacEwan University.

---

**Supplementary Table 1.**
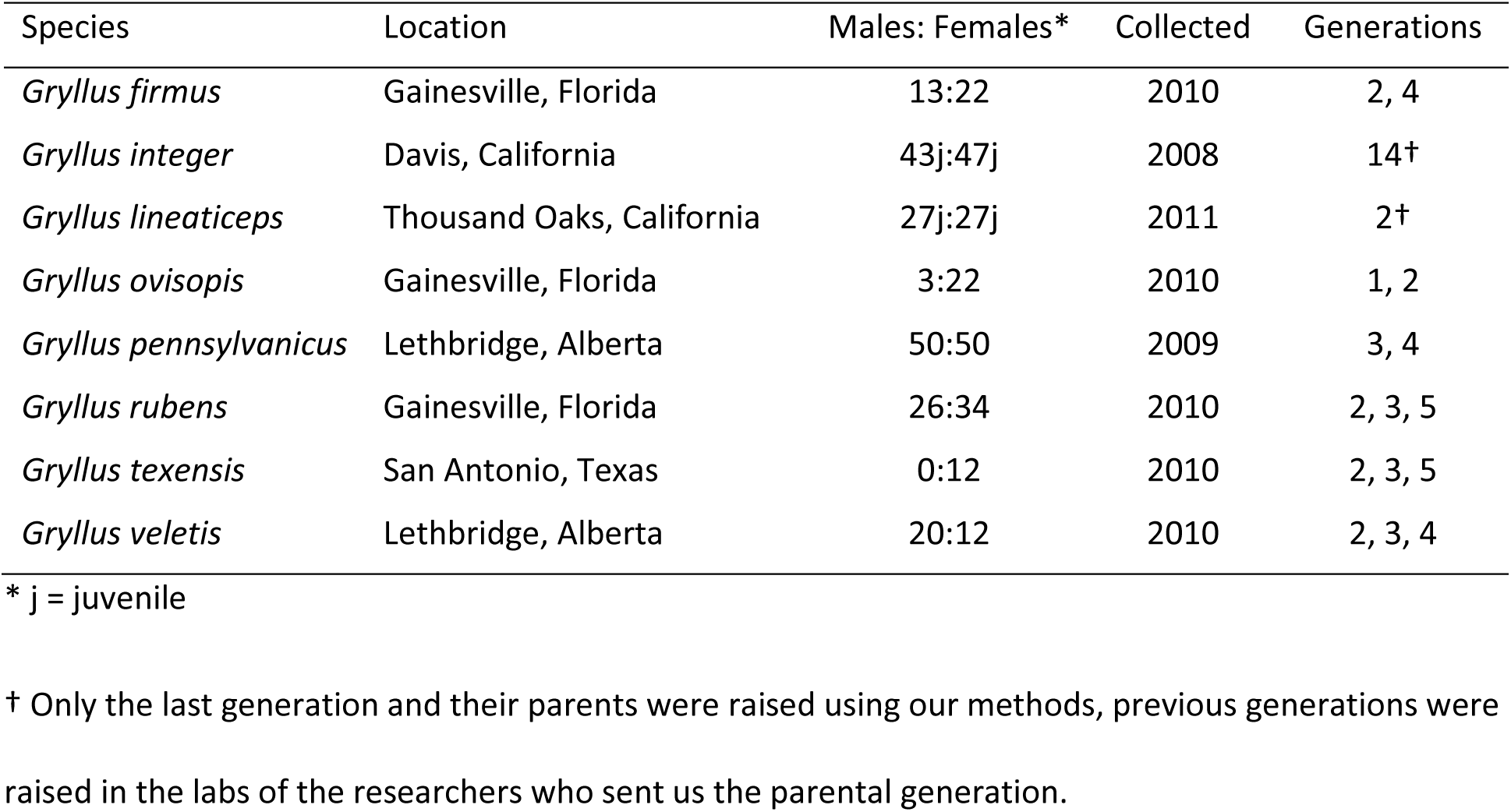
Sources of field crickets used in this study, the number and sex ratio of the parental generation of our lab colonies, the year they were collected from the wild, and the filial generations used in this study.

