## Supplementary Table 1 for "The evolution of weaponry and aggressive behaviour in field crickets"

Supplementary Table 1. Sources of field crickets used in this study, the number and sex ratio of the parental generation of our lab colonies, the year they were collected from the wild, and the filial generations used in this study.

| Species | Location | Males: Females* | Collected | Generations |
|---|---|---|---|---|
| *Gryllus firmus* | Gainesville, Florida | 13:22 | 2010 | 2, 4 |
| *Gryllus integer* | Davis, California | 43j:47j | 2008 | 14† |
| *Gryllus lineaticeps* | Thousand Oaks, California | 27j:27j | 2011 | 2† |
| *Gryllus ovisopis* | Gainesville, Florida | 3:22 | 2010 | 1, 2 |
| *Gryllus pennsylvanicus* | Lethbridge, Alberta | 50:50 | 2009 | 3, 4 |
| *Gryllus rubens* | Gainesville, Florida | 26:34 | 2010 | 2, 3, 5 |
| *Gryllus texensis* | San Antonio, Texas | 0:12 | 2010 | 2, 3, 5 |
| *Gryllus veletis* | Lethbridge, Alberta | 20:12 | 2010 | 2, 3, 4 |

* j = juvenile

† Only the last generation and their parents were raised using our methods, previous generations were raised in the labs of the researchers who sent us the parental generation.
